# Audiovisual Integration in the Human Brain: A Coordinate-based Meta-analysis

**DOI:** 10.1101/2022.07.13.499939

**Authors:** Chuanji Gao, Jessica J. Green, Xuan Yang, Sewon Oh, Jongwan Kim, Svetlana V. Shinkareva

## Abstract

People can seamlessly integrate a vast array of information from what we see and hear in the noisy and uncertain world. However, the neural underpinnings of audiovisual integration continue to be the topic of debate. Using strict inclusion criteria, we performed activation likelihood estimation meta-analysis on 121 neuroimaging experiments with a total of 2,092 participants. We found that audiovisual integration is linked with the coexistence of multiple integration sites including early cortical, subcortical, and higher association areas. Although activity was consistently found within the superior temporal cortex, different portions of this cortical region were identified depending on the analytical contrast used, complexity of the stimuli, and modality within which attention was directed. The context-dependent neural activity related to audiovisual integration suggests a flexible rather than fixed neural pathway for audiovisual integration. Together, our findings highlight a flexible multiple pathways model for audiovisual integration, with superior temporal cortex as the central node in these neural assemblies.

---

Much of the information we encounter in the environment is noisy and ambiguous. Integrating information from multiple sensory systems allows us to make better inferences about what we are experiencing (Ernst & Bülthoff, 2004; Noppeney, 2021). For instance, in a busy restaurant it may be difficult to hear what your friend is saying, but by integrating the speech sounds with mouth movements can greatly increase your understanding of the conversation. Although many models with good fit for behavioral indices of audiovisual integration have been developed to explain how information from multiple senses is combined (Colonius, 1990; Colonius & Diederich, 2020; Deneve & Pouget, 2004; Ernst & Banks, 2002; Fetsch et al., 2012; Körding et al., 2007; Magnotti et al., 2013; Miller, 1982; Parise & Ernst, 2016; Shams & Beierholm, 2010; Shams et al., 2005), it remains unclear how the efficient combination of audiovisual information is accomplished in the brain. Here, we used a meta-analytic approach to identify common patterns of brain activity across a wide variety of audiovisual studies.

Different theories on the neural basis of audiovisual integration have been put forth (see Driver & Noesselt, 2008; Stein & Stanford, 2008 for reviews). In line with a traditional hierarchical route, the *Higher Association Areas* model states that unisensory signals are processed independently in their respective sensory cortices and integration occurs in later association areas such as superior temporal cortex [**Fig 1**]. However, accumulating evidence indicates that integration might occur at sensory-perceptual [Sensory areas model] and subcortical levels [Subcortical areas model] prior to any processing in association cortices [**Fig 1**]. Though these models have been around for many years, it is still unclear which model accurately reflects the neural substrates of audiovisual integration. This lack of consensus across studies could be due to a number of factors, including analytical contrasts, stimulus complexity and attention. There is variability in how audiovisual neural activity is defined and analyzed (Calvert & Thesen, 2004; Stein et al., 2010), as well as a wide variety of stimulus types and attentional manipulations used that could influence the patterns of observed brain activities. How much does the neural activity of audiovisual integration overlap across these different factors? If the neural activity is essentially the same irrespective of these factors, it would suggest that audiovisual integration is implemented by a fixed network of brain regions. If, however, audiovisual integration is implemented by different brain regions depending on experimental and analytical contexts, it would suggest that the neural pathways involved are more flexible and there may not be a single neural network responsible for integration.

**Fig 1.**
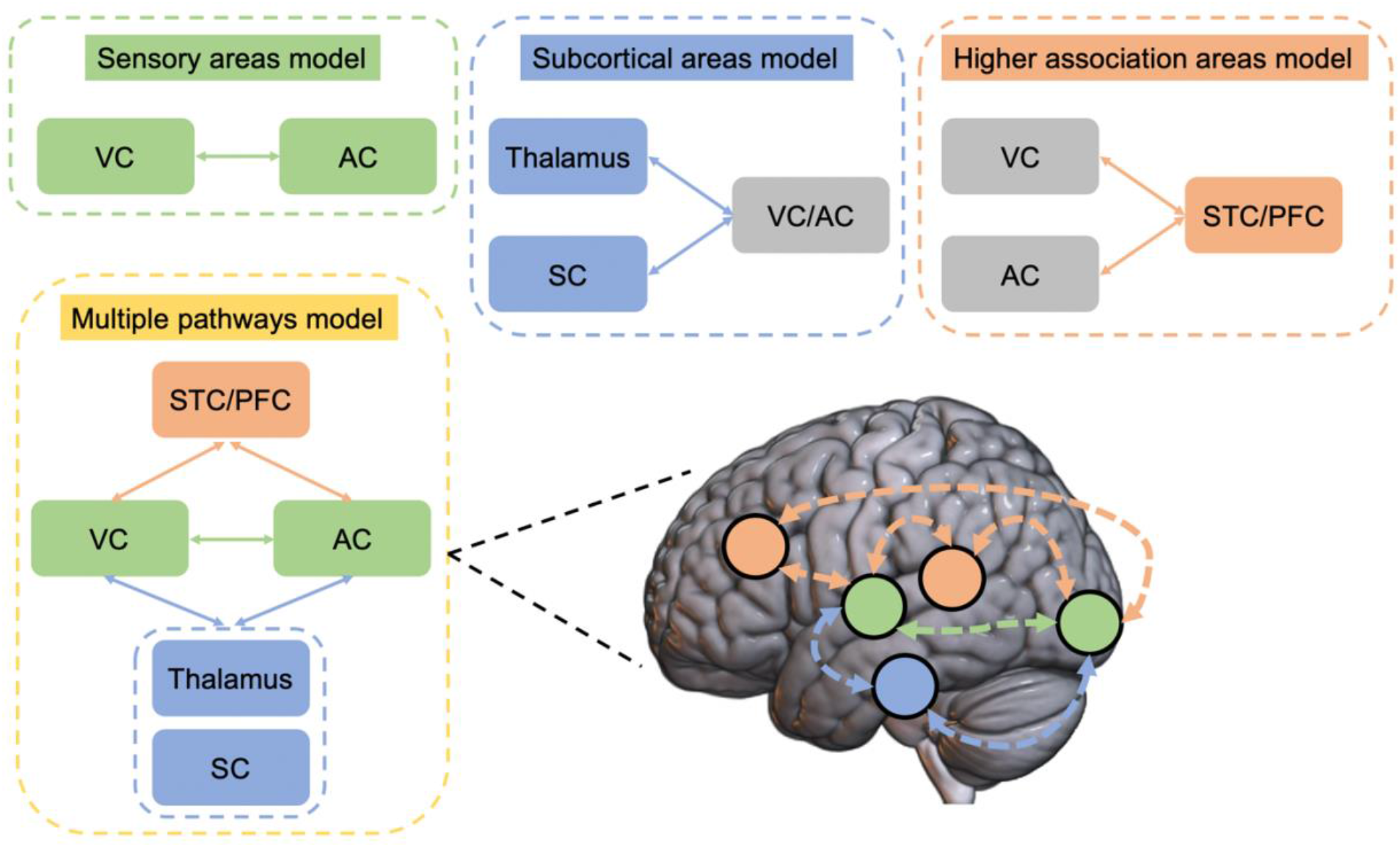
Possible neural pathways for audiovisual integration. Sensory areas model posits that direct connections between visual cortex (VC) and auditory cortex (AC) are underlying the audiovisual integration process. Subcortical areas model posits that audiovisual integration takes place at subcortical areas such as thalamus or superior colliculus (SC). High association areas model posits that audiovisual integration takes place at higher association areas such as superior temporal cortex (STC) and prefrontal regions (PFC). Multiple pathways model posits coexistence of all the pathways, in which sensory areas, subcortical areas, and high association areas are all related to this process.

There are three main approaches widely used in examining the neural activity of audiovisual integration. The classic approach to examining audiovisual integration is to compare neural responses to an audiovisual stimulus with the sum of the unisensory responses (i.e., AV > A+V), which is herein referred to as the “Interaction” approach. This approach assumes that audiovisual integration results in an interaction between modalities that is more than the sum of the unisensory parts. Such a pattern has been clearly identified at the cellular level (Meredith et al., 1987; Meredith & Stein, 1983, 1986a, 1986b; Stein & Stanford, 2008; Wallace et al., 1996) and this interaction approach has subsequently been used to examine audiovisual integration non-invasively in humans using a number of techniques and paradigms. For example, Calvert et al. (2000) presented subjects with audiovisual speech signals and each modality in isolation. Using this interaction contrast, they found that left superior temporal sulcus exhibited significant supra-additive responses, where the response to the audiovisual speech was greater than the summed unimodal responses. However, a number of limitations to this comparison have been identified, such as being overly conservative when used in neuroimaging studies due to ‘hemodynamic refractoriness’ (Friston et al., 1998; Mechelli et al., 2001), inappropriately double-subtracting activity that is common to both auditory and visual tasks when the unimodal stimuli are summed together (Teder-Sälejärvi et al., 2002), and only producing a reliable interaction effect for weak or sub-threshold stimuli (Holmes, 2007; Meredith & Stein, 1983).

In an attempt to overcome some of the issues with the interaction approach, many studies have used alternative analytical contrasts to identify brain activity associated with audiovisual integration. One common approach, herein referred to as the “Conjunction” approach, compares the audiovisual response to each unimodal response and then identifies overlap between the two [i.e., (AV > V) ∩ (AV > A)]. This approach is based on similar electrophysiological findings as the “Interaction” approach and is presumably less conservative. For example, ENREF 2 ENREF 4 one study measured fMRI signals during presentations of pictures of animals and other objects, environmental sounds and audiovisual clips. They found overlapping activation in a series of brain areas such as thalamus, inferior occipital gyrus, and lingual gyrus, that have stronger responses to audiovisual stimuli than either unimodal condition (Belardinelli et al., 2004). One potential limitation of the Conjunction approach is that it can reflect common brain activations across unimodal visual and auditory stimuli but not audiovisual integration per se, in which (AV > V) ∩ (AV > A) would be equal to V ∩ A (Calvert & Thesen, 2004; Ethofer et al., 2006).

A third common approach focuses on comparing neural responses for congruent and incongruent audiovisual stimuli, herein referred to as the “Congruency” approach [Congruent > Incongruent]. This comparison assumes that the congruency of the stimuli can lead to different magnitudes of integration (Kadunce et al., 1997; Wallace et al., 1996). For example, one study instructed participants to attend to lip movements that were either congruent or incongruent with spoken words ENREF 67. They found increased activation in the superior temporal cortex and other brain areas for congruent compared to incongruent conditions (Fairhall & Macaluso, 2009). One limitation of this approach is that it can identify brain activations that are related to audiovisual interactions (i.e., modulation effects from one modality on the other) but not necessarily integration.

In addition to variability in how audiovisual brain activities are analyzed, stimuli in studies of audiovisual integration vary in terms of complexity from very simple stimuli (e.g., flashes and tones) to complex stimuli (e.g., object images and sounds or speech stimuli). Compared to simple stimuli, complex stimuli may involve integration of both low-level spatial and temporal (Stein & Stanford, 2008) and higher-level semantic information (Doehrmann & Naumer, 2008). Though many studies suggest a common cognitive process for audiovisual integration irrespective of different stimulus types (Driver & Noesselt, 2008; Hocking & Price, 2008; Stekelenburg & Vroomen, 2007; Stevenson & James, 2009), there is evidence that suggests the opposite (Tuomainen et al., 2005). It is not yet clear whether the neural mechanisms of audiovisual integration would differ depending on different types of stimuli.

Lastly, brain regions engaged by audiovisual integration could also depend on attention. Previous literature has demonstrated the role of top-down attention in modulating audiovisual integration (Macaluso et al., 2016; Noppeney, 2021; Rohe & Noppeney, 2018; Talsma, 2015; Talsma et al., 2010). For example, Talsma et al. (2006) observed a supra-additive effect on neural activity (as measured with event-related potentials) when the stimuli were attended, but this effect reversed when the stimuli were unattended. Across neuroimaging studies of audiovisual integration, a wide variety of tasks have been used that vary in how attention is directed to the stimuli. Some tasks require attention to only one of the sensory modalities while the other is ignored (e.g., Bonath et al., 2007; Green et al., 2009; Noppeney et al., 2010), some require attention to both modalities (e.g., Benoit et al., 2010; Bushara et al., 2001; Hocking & Price, 2008), and still others use passive tasks in which none of the stimuli are task-relevant (e.g., Barrós-Loscertales et al., 2013; Butler et al., 2011; Holle et al., 2010). With the accumulating evidence that audiovisual integration can be heavily dependent on the attended modality (Talsma, 2015), it seems likely that this variability in attention demands across tasks could modify what brain activity is observed.

The present study used a meta-analysis approach with the activation likelihood estimation (ALE) method (Eickhoff et al., 2012; Eickhoff et al., 2009; Turkeltaub et al., 2012) to identify the brain areas associated with audiovisual integration. First, by aggregating neuroimaging studies of audiovisual integration, this meta-analysis study allowed us to identify brain regions commonly engaged by audiovisual integration while overcoming many of the limitations of individual studies, such as small samples, lack of confirmation for isolated findings, and generalization of context-specific findings. Second, we examined whether there are distinct patterns of activation associated with varying analytical contrasts, stimulus complexities, and attentional demands. By identifying shared and unique voxels across conditions, these analyses enabled us to disentangle areas that have a central role in integration, independent of context, from brain areas that show context-dependent integration activity.

## Materials and Methods

### Study identification

Literature search was conducted across PubMed and Web of Science databases. Search terms included combinations of the three descriptors: one *mandatory research question descriptor* (“integration”), one *research question descriptor* (e.g., “multisensory”, “audiovisual” etc.) and one *methodological descriptor* (e.g., “fMRI”, “MRI”, “PET”, “BOLD”) for studies published until September 2020 [See **Supplemental info** for specific search terms used in study identification]. In PubMed, the search was performed within the title and abstract, with preliminary filters including 1) papers written in English; 2) papers published in journal articles; 3) human adult subjects. In Web of Science, the search was also performed within title and abstract, with preliminary filters including 1) papers written in English; 2) papers published in journal articles; and 3) not a study using animal subjects. The overall search revealed a total of 1,118 studies. After removing duplicated records, 741 articles remained. A step-by-step flowchart of the meta-analysis procedure is shown in **Fig S1**. The meta-analysis was performed with PRISMA standards (Liberati et al., 2009).

### Study screening

We then assessed the full-text of the 741 articles for eligibility with the following criteria: 1) It reported original data; 2) It is a neuroimaging study using PET or fMRI; 3) It included healthy subjects; 4) It included adult subjects (≥ 18 years old); 5) It included whole brain statistics; 6) It included coordinates reported in Talairach or MNI space; 7) It included a relevant contrast. To be considered a relevant contrast, the contrast had to involve a multisensory condition and had to be interpretable as multisensory integration and not unimodal sensory differences. For example, the conjunction of visual only and auditory only conditions was excluded because without an audiovisual comparison it cannot speak directly to multisensory integration, and a contrast of audiovisual > visual was excluded because the results could be interpreted as differences in auditory sensory processing.

After the study screening, a total of 98 studies were included. An additional 39 articles were included by tracing the articles cited by the included studies and reading relevant review articles. Thus, the final dataset included 137 papers that reported 139 experiments. The articles were divided into eight groups that were assessed by four independent researchers (CG, JK, SO, and XY). Another independent check of the screening results was also performed, in which the assignment was then switched between the four researchers. For articles where there was initial disagreement, consensus was reached through discussion among all authors.

### Data extraction

For each study, we extracted the following data: 1) study ID (first author, publication year, and journal); 2) imaging modality (MRI or PET); 3) MRI strength (e.g., 3T); 4) sample size; 5) sensory modality (visual-auditory and visual-tactile); 6) stimuli; 7) task; 8) contrast; 9) normalization space (MNI or Talairach); 10) peak coordinates (x/y/z); and 11) analysis package [**Supplemental info, Datasets S1-2**].

### Statistical methods

#### Activation likelihood estimation

Coordinate-based meta-analyses were conducted with the revised ALE algorithm, implemented in GingerALE 3.0.2 (BrainMap, http://brainmap.org/ale/). Coordinates reported in Talairach space were transformed into MNI coordinates using a linear transformation to perform analyses in a common stereotactic space (Lancaster et al., 2007). In the ALE analyses, each reported location was taken as the center of a 3D Gaussian probability density distribution. The uncertainty associated with localization of each location was modelled by the full-width at half-maximum of the Gaussian function determined by the number of participants in each study. After that, a modelled activation (MA) map was created for each voxel reflecting the probability of an activation at that location. A 3D ALE map was created by taking the union across all of the MA maps. The voxel-specific ALE scores reflect the consistency of the activation locations. An empirically derived null-distribution was achieved by sampling a voxel at random from each of the MA maps and taking the union of these values in the same manner as the true analysis. A voxel-wise *p*-map was then created by comparing the ALE scores to the null-distribution, which was then submitted to a cluster-level family-wise error (FWE) correction with a cluster level threshold of *p* < .05 and a cluster-forming threshold at voxel-level *p* < .001. The significant cluster size was determined by comparison to a null-distribution of cluster-sizes derived by simulating 1,000 datasets of randomly distributed foci with a threshold of *p* < .05. Anatomical areas were labeled using the SPM Anatomy Toolbox. Results were visualized using MRIcroGL (https://www.nitrc.org/projects/mricrogl/) for slices and FreeSurfer (https://surfer.nmr.mgh.harvard.edu/) for surface rendering. A nonlinear mapping tool was used to convert volumetric to surface coordinates for display purposes (Wu et al., 2018).

### Experiment categorization and analyses

Based on the purpose of the present study, we classified the included experiments in the following way: 1) Sensory modality: visual-auditory or visual-tactile. 2) Contrast, for the two modalities (unimodal1 and unimodal2): (multisensory congruent > unimodal1) ∩ (multisensory congruent > unimodal2); (multisensory congruent < unimodal1) ∩ (multisensory congruent < unimodal2); (multisensory incongruent > unimodal1) ∩ (multisensory incongruent > unimodal2); multisensory congruent > sum of unimodal; multisensory congruent < sum of unimodal; multisensory congruent > max(unimodal1, unimodal2); multisensory incongruent < max(unimodal1, unimodal2); multisensory congruent > mean(unimodal1, unimodal2); integration > no integration; no integration > integration; congruent > incongruent; incongruent > congruent; illusion > no illusion; and no illusion > illusion. 3) Stimuli: simple, complex and speech. Here the stimuli were coded as simple, such as flashes and pure sounds; complex, such as pictures of objects, sounds of objects; and speech, such as speech videos and speech sounds. 4) Attention: audiovisual, visual, auditory, auditory or visual or audiovisual, auditory and visual, visuotactile, tactile, visual or tactile or visuotactile, and passive. Note that ‘auditory or visual or audiovisual’ means that the contrast involves both multisensory and unimodal conditions, in which participants were asked to pay attention to both modalities for multisensory condition, and they were asked to pay attention to either of the single modality for unimodal conditions. 5) Illusion: illusion and no illusion.

We performed the following analyses:

1.1. Sensory modality: visual-auditory, 121 experiments.
1.2. Sensory modality: visual-tactile, 18 experiments.
2.1. Contrast: (multisensory congruent > unimodal1) ∩ (multisensory congruent > unimodal2), 28 experiments.
2.2. Contrast: multisensory congruent > sum of unimodal, 23 experiments.
2.3. Contrast: congruent > incongruent, 44 experiments.
2.4. Contrast: incongruent > congruent, 39 experiments.
3.1. Stimuli: simple, 24 experiments.
3.2. Stimuli: complex-nonspeech, 23 experiments.
3.3. Stimuli: complex-speech, 75 experiments.
4.1. Attention: audiovisual, 59 experiments. Note that we combined “audiovisual” (29 experiments) and “auditory or visual or audiovisual” (30 experiments) in one category because audiovisual modality was attended in the audiovisual conditions for the experiments of “auditory or visual or audiovisual”.
4.2. Attention: visual, 14 experiments.
4.3. Attention: auditory, 19 experiments.
4.4. Attention: passive, 27 experiments.

Given the small number of experiments for illusion studies (16 experiments), we did not perform analyses for illusion and no illusion conditions separately due to a higher risk that results can be driven by single experiments for a small sample size (Eickhoff et al., 2016). A series of other conditions [e.g., (multisensory congruent < unimodal1) ∩ (multisensory congruent < unimodal2)] were also not included for analyses for the same reason.

To determine the overlap in neural activity across conditions, we performed conjunction analyses using the conservative minimum statistic, in which only brain voxels significant on a corrected level in each individual analysis were considered (Nichols et al., 2005).

## Results

Our study screening procedures identified 139 neuroimaging experiments involving healthy adult participants that directly assessed multisensory integration in their analyses and provided whole-brain results. Of these studies, 121 examined audiovisual integration while only 18 examined visual-tactile integration and no studies meeting our criteria examined tactile-auditory integration. Because of this bias in the literature, we were unable to address our original question of whether or not there is a general multisensory integration network. Furthermore, because of the small number of visual-tactile studies, further categorizing those studies based on stimulus or task features would increase the risk that the results would be driven by a single experiment (Eickhoff et al., 2016). Thus, we restricted our subsequent analyses to the 121 audiovisual studies. However, all the visual-tactile studies are included in [**Supplemental info, Table S1**] for completeness.

### Audiovisual Integration is Linked with Multiple Integration Sites including Early Sensory, Subcortical and Higher Association Areas

The meta-analysis of audiovisual integration showed convergence in bilateral superior temporal gyrus, left middle temporal gyrus and fusiform gyrus, left middle temporal gyrus, bilateral middle occipital gyrus, bilateral inferior occipital gyrus, right inferior frontal gyrus, bilateral middle frontal gyrus, left precentral gyrus, bilateral insula, right thalamus, left medial frontal gyrus, left superior frontal gyrus, and right lingual gyrus [**Supplemental info, Fig S2 and Table S1**]. These findings suggest that audiovisual integration is associated with multiple integration sites including early cortical areas, subcortical areas, and higher association areas.

### Brain Regions Engaged by Audiovisual Integration Depend on Analytical Contrast, Stimulus Complexity, and Attention

We further analyzed these audiovisual studies by categorizing the experiments based on key factors that we hypothesized could influence how audiovisual integration is observed in the brain: the contrast used for comparison, the complexity of the stimuli, and how participants’ attention was directed during the task.

#### Brain Regions Engaged by Audiovisual Integration Depend on Analytical Contrast

To explore how the contrasts used to define audiovisual integration influence the brain regions identified, we separated studies into three broad categories: (1) “Conjunction”: contrasts that focus on the overlap between the audiovisual response and each unimodal response [i.e., (AV > V) ∩ (AV > A); n=28]; (2) “Interaction”: contrasts that focus on supra-additive responses, where the audiovisual response is greater than the sum of the unimodal responses [i.e., AV > A+V; n=23]; and 3) “Congruency”: contrasts that focus on comparing congruent and incongruent audiovisual stimuli. We further separated these contrasts by whether they were focused on the congruent stimuli [Congruent > Incongruent, n=44] or the incongruent stimuli [Incongruent > Congruent, n=39], as these would highlight different cognitive and perceptual processes.

The “Conjunction” contrast refers to (AV > V) ∩ (AV > A). The “Interaction” contrast refers to AV > A+V. The “Congruency” contrast refers to “Congruent > Incongruent”. The legend indicates the number of significant voxels identified for each contrast and their conjunctions. Surface rendering was created using converted surface coordinates from MNI coordinates for visualization purpose.

The meta-analysis of Conjunction contrast showed activation in left and right superior temporal gyrus, left and right thalamus, and right parahippocampal gyrus. The meta-analysis of Interaction contrast showed activation in left and right superior temporal gyrus, left middle temporal gyrus, and right postcentral gyrus. The meta-analysis of Congruent > Incongruent contrast showed activation in left and right superior temporal gyrus, left inferior temporal gyrus, left inferior occipital gyrus, and left middle occipital gyrus. The meta-analysis of Incongruent > Congruent contrast showed activation in right superior temporal gyrus, left middle frontal gyrus, left superior frontal gyrus, right inferior frontal gyrus, and left middle temporal gyrus [**Fig 2; Supplemental info, Table S2**]. These findings showed that brain regions engaged by audiovisual integration varied to a great extent depending on different analytical contrasts.

**Fig 2.**
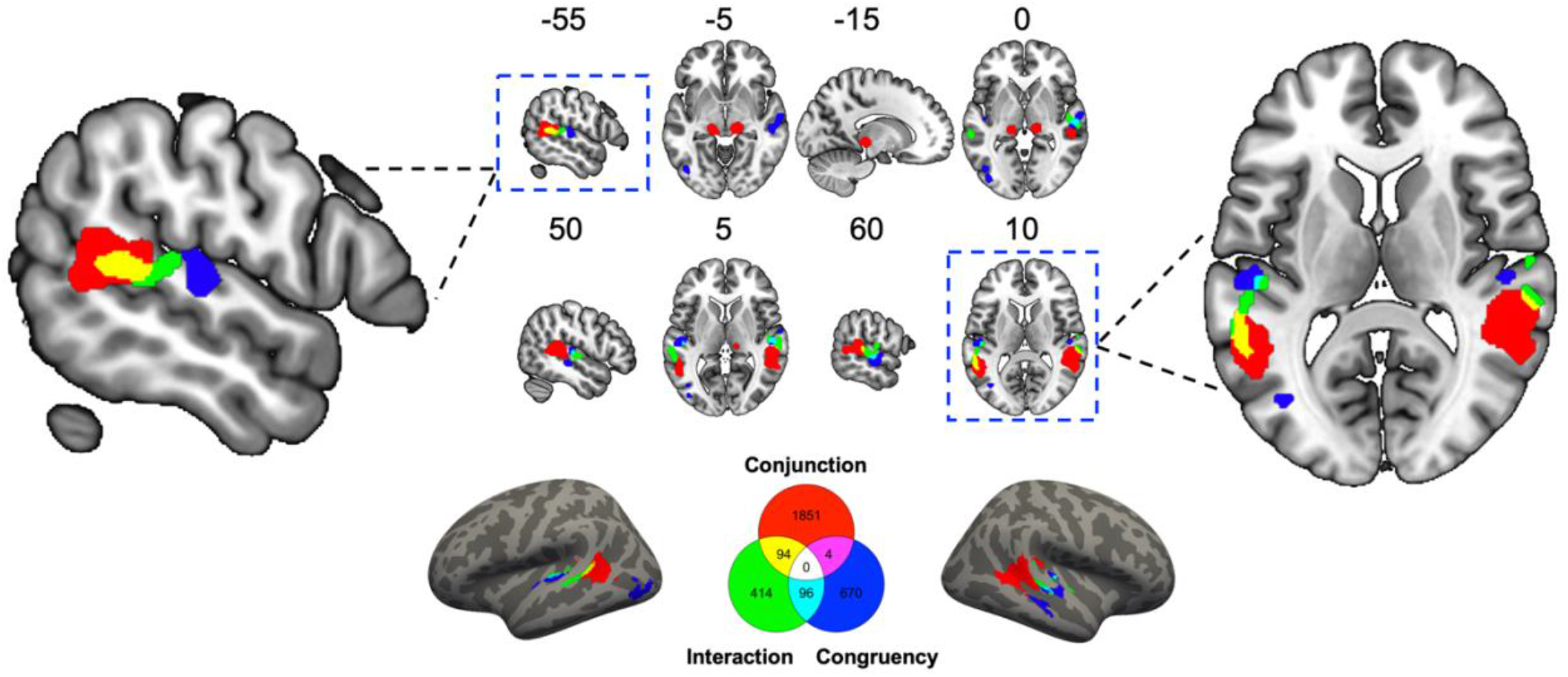
Significant brain regions from the activation likelihood estimation (ALE) meta-analyses and number of shared and unique voxels shown by a Venn diagram for effects of analytical contrast. These findings showed that brain regions linked with audiovisual integration varied to a great extent depending on different analytical contrasts. Shared voxels across contrasts were mainly located at superior temporal cortex [see also **Table S3**], suggesting its central role in audiovisual integration.

The common activation between Conjunction and Interaction contrasts included right superior temporal and left middle temporal cortices. The common activation between Conjunction and Congruency contrasts included right superior temporal cortex. The common activation between Interaction and Congruency [Congruent > Incongruent] contrasts included bilateral superior temporal cortices [**Fig 2; Supplemental info, Table S3**]. These analyses showed that the pairwise overlapped brain voxels across analytical contrasts were consistently located within the superior temporal cortex, although no voxels were commonly activated by all three contrasts.

#### Brain Regions Engaged by Audiovisual Integration Depend on Stimulus Complexity

We then compared how stimulus complexity influences audiovisual integration. As some studies have suggested there may be differences in how speech is processed compared to other complex non-speech stimuli, we categorized studies into those using Simple stimuli [e.g., flashes and beeps; n=24], Complex-nonspeech [e.g., pictures and sounds of objects; n=23], and Complex-speech [e.g., spoken phonemes or words along with visible mouth movements; n=75].

The meta-analysis of Simple stimuli showed activation in bilateral superior temporal gyrus, right transverse temporal gyrus, right middle and inferior frontal gyrus. The meta-analysis of Complex-nonspeech stimuli showed activation in right superior temporal gyrus. The meta-analysis of Complex-speech stimuli showed activation in bilateral superior temporal gyrus, left middle temporal gyrus, left superior, medial, middle and inferior frontal gyrus, right middle and inferior occipital gyrus, right lingual gyrus, and right thalamus [**Fig 3; Supplemental info, Table S4**]. These findings showed that brain regions engaged by audiovisual integration varied to a great extent depending on different stimulus complexity.

**Fig 3.**
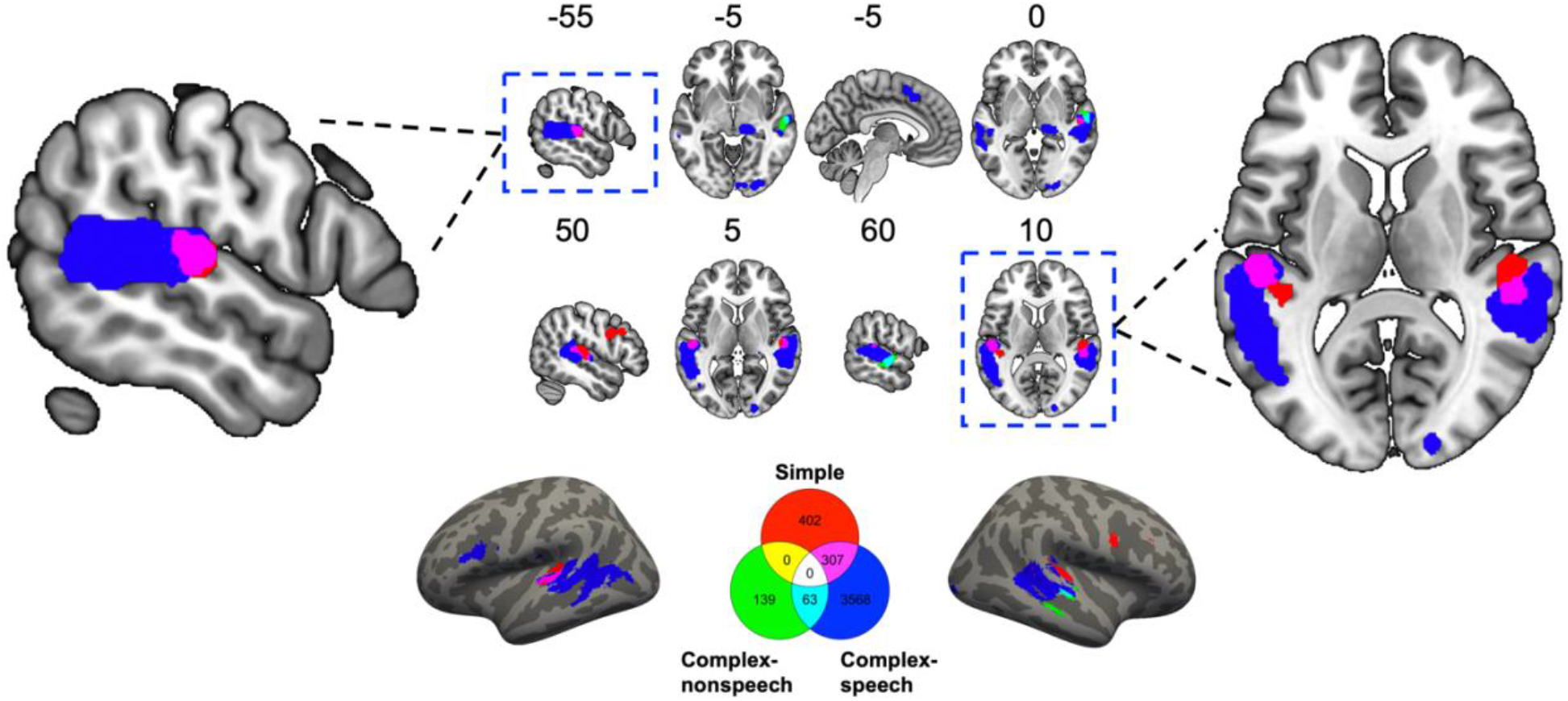
Significant brain regions from the activation likelihood estimation (ALE) meta-analyses and number of shared and unique voxels shown by a Venn diagram for effects of stimulus complexity. These findings showed that brain regions linked with audiovisual integration varied to a great extent depending on different stimulus complexity. Shared voxels across contrasts were mainly located at superior temporal cortex [see also **Table S5**], suggesting its central role in audiovisual integration. The legend indicates the number of significant voxels identified for each contrast and their conjunctions. Surface rendering was created using converted surface coordinates from MNI coordinates for visualization purpose.

The common activation between Simple and Complex-speech included bilateral superior temporal cortices. The common activation between Complex-nonspeech and Complex-speech included right superior temporal cortex [**Fig 3; Supplemental info, Table S5**]. These analyses showed that the pairwise overlapped brain voxels across different stimulus complexity were consistently located within the superior temporal cortex, although, as with the comparison of analytical contrasts, no voxels were commonly activated by all three stimulus types.

#### Brain Regions Engaged by Audiovisual Integration Depend on Attention

Lastly, we compared how the modality that participants were instructed to attend to influences the integration activations that are observed. To do this we categorized studies into those that had participants attend to the audiovisual stimuli (n=59; this includes studies that have participants attend to both modalities when present, but may have them attend to only one modality when it is the only one presented), those that had participants only attend to the visual information (n=14), just the auditory information (n=19), and those in which participants did not actively attend to any of the stimuli (i.e., passive tasks; n=27).

The meta-analysis of Audiovisual attention showed activation in right insula, right middle temporal gyrus, and right superior temporal gyrus. The meta-analysis of Visual attention showed activation in right superior temporal gyrus. The meta-analysis of Auditory attention showed activation in left superior temporal gyrus, left medial frontal gyrus, left and right insula. The meta-analysis of Passive attention showed activation in bilateral superior temporal gyrus, bilateral middle temporal gyrus, left middle frontal gyrus, right middle occipital gyrus, and right lingual gyrus [**Fig 4; Supplemental info, Table S6**]. These findings showed that brain regions engaged by audiovisual integration varied to a great extent depending on different attention conditions.

**Fig 4.**
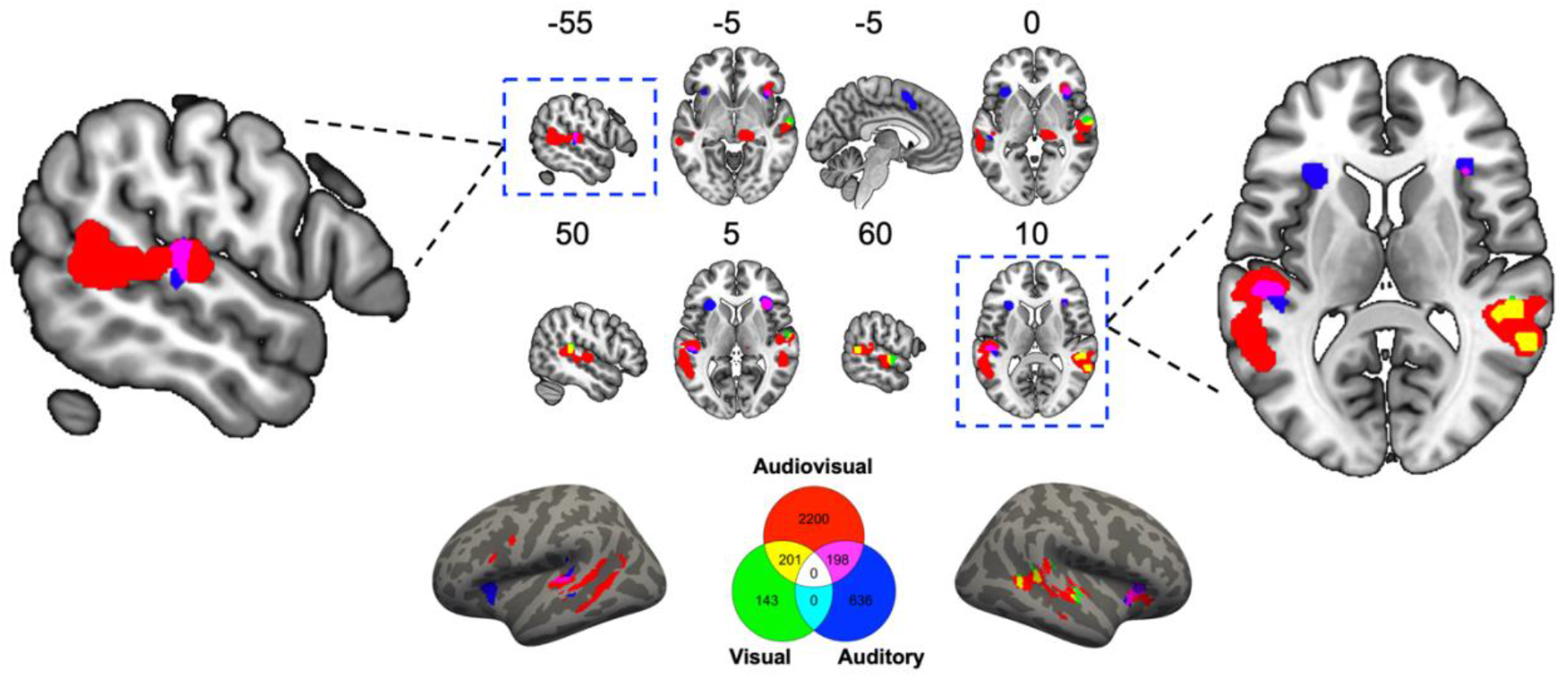
Significant brain regions from the activation likelihood estimation (ALE) meta-analyses and number of shared and unique voxels shown by a Venn diagram for effects of attention. These findings showed that brain regions linked with audiovisual integration varied to a great extent depending on different attention conditions. Shared voxels across contrasts were mainly located at superior temporal cortex [see also **Table S7**], suggesting its central role in audiovisual integration. The legend indicates the number of significant voxels identified for each contrast and their conjunctions. Surface rendering was created using converted surface coordinates from MNI coordinates for visualization purpose.

The common activation between Audiovisual attention and Visual attention included right superior temporal cortex. The common activation between Audiovisual attention and Auditory attention included right insula and left superior temporal cortex [**Fig 4; Supplemental info, Table S7**]. As with the previous analyses, the pairwise overlapped brain voxels across different attention conditions were consistently located within superior temporal cortex, but there were no voxels commonly activated across the attention conditions.

## Discussion

What are the brain regions associated with audiovisual integration? This question has been examined for many years without reaching a consensus. Here, we summarized 121 neuroimaging experiments comprised of 2092 participants, all of which examined the neural basis of audiovisual integration with strict inclusion criteria. Moreover, we examined how experimental context and analytical choices influence the brain networks identified during audiovisual integration, leading to three main findings. First, audiovisual integration is associated with multiple integration sites including early cortical areas, subcortical areas, and higher association areas, which is consistent with a multiple pathways model. Second, neural assemblies of audiovisual integration varied to a great extent depending on analytical contrast, stimulus complexity and attention, suggesting a flexible rather than a fixed neural pathway model for audiovisual integration. Third, neural activity consistently occurred at superior temporal cortex, albeit in slightly different locations based on context, suggesting its central role in audiovisual integration.

Our results suggest that audiovisual integration occurs at multiple levels: sensory sites including middle and inferior occipital gyrus, fusiform gyrus and lingual gyrus, and middle portion of superior temporal gyrus; subcortical sites including thalamus; higher association sites including superior temporal cortex, middle and superior frontal gyrus. Though the evidence is correlational rather than causal, our findings are consistent with a multiple pathways model as outlined in **Fig 1**, in which audiovisual integration is not restricted to a particular brain area, but can occur via engaging a network of brain regions. Based on this model, audiovisual integration may partly occur via direct communication between visual and auditory cortices, in which traditional unisensory regions may have multisensory characteristics (Brosch et al., 2005; Calvert et al., 1997; Cappe & Barone, 2005; Falchier et al., 2002; Foxe et al., 2000; Ghazanfar & Schroeder, 2006; Mishra et al., 2007; Rockland & Ojima, 2003; Shams et al., 2001). There may also be a cortical-thalamic-cortical route in which sensory information is sent from sensory cortices, integrated at thalamus, and sent back to a cortical region such as prefrontal cortex (Cappe et al., 2009; Cappe et al., 2009; Hackett et al., 2007). This evidence suggests that the thalamus may have a function of computation (i.e., integration) beyond its role of relaying information (Rikhye et al., 2018). Higher association cortices including superior temporal cortex and prefrontal cortex have been shown to connect with visual and auditory sensory cortices (Fuster et al., 2000; Romanski et al., 1999; Seltzer & Pandya, 1994), in which there are bidirectional feedforward and feedback communications among these brain regions. Although it is possible that our overall results were influenced by differences in the number of studies across conditions, our separate analyses of each condition with conjunction analyses to identify common loci of activity mitigates this limitation. The results were not driven by a specific type of task, as task types are diverse [**Supplemental info, Dataset S1**]. Thus, our results support the idea that audiovisual integration can take place at a myriad of levels throughout the cortex.

Audiovisual integration appears to be context-dependent. Analytical contrasts (i.e., criteria) for defining neural activity as audiovisual integration are diverse and under debate. This issue is further complicated by different stimulus complexity (simple, complex-nonspeech or complex-speech) and attention (attending audiovisual, visual or auditory). Our analyses revealed that there were no shared brain voxels across different conditions for either analytical contrasts, stimulus complexity, or attention, and there were only a small portion of shared voxels for each pairwise comparison. These findings suggest that the neural correlates of audiovisual integration are highly context dependent, consistent with previous evidence that audiovisual integration is flexible and context dependent (see Van Atteveldt et al., 2014 for a review).

Our results showed that the ‘Conjunction’ contrast had the largest number of activated voxels and the ‘Interaction’ contrast had the smallest number of activated voxels, which is consistent with previous literature suggesting that the ‘Interaction’ contrast is a very strict criterion that will likely lead to missed activity (Beauchamp, 2005). It is possible that the ‘Interaction’ contrast, which was adopted from single-neuron studies examining changes in action potentials, does not accurately capture the integration-related changes in BOLD activity, which predominantly reflects postsynaptic activity (Laurienti et al., 2005; Stanford & Stein, 2007). This difference in sensitivity of the contrasts might explain why a subcortical node (i.e., thalamus) was only identified for the ‘Conjunction’ contrast. In addition to the thalamus, differences between the analytical contrasts appeared in other regions as well. We observed an anterior-posterior differentiation in the superior temporal cortex across contrasts, with the congruency contrast activity being located more anteriorly compared to the interaction or conjunction contrast activity. It is possible that the congruency contrast reveals modulations of one modality on another modality in addition to the integration of inputs. In line with this interpretation, we also found the involvement of visual regions such as occipital lobe and inferior temporal gyrus only for the congruency contrast. Additionally, the inferior frontal cortex was only identified for the ‘Incongruent > Congruent’ contrast. It is likely that the inferior frontal cortex activation reflects conflict resolution rather than an integration-specific process, given previous evidence on the link between inferior frontal cortex and domain-general conflict resolution and cognitive control (Derrfuss et al., 2005; Nelson et al., 2009).

We also found an anterior-posterior distinction between simple/complex-non-speech (anterior) and complex speech stimuli (posterior) in the superior temporal cortex. Posterior superior temporal cortex (traditionally labelled as Wernicke’s area) has been associated with speech perception and comprehension (Lesser et al., 1986; Naeser et al., 1987; But see also Binder, 2017). However, the middle portions of superior temporal cortex has been linked with processing of auditory inputs or objects (Price, 2010). Thus, one interpretation of our findings is that brain regions relevant for processing particular stimulus types are directly involved in the audiovisual integration of those specific stimuli. Consistent with this interpretation, we only found activation of left inferior frontal gyrus (another classical language area, Broca’s area) for audiovisual integration of speech but not for simple or complex-non-speech stimuli. Moreover, there were more activated voxels in the left than the right hemisphere for audiovisual integration of speech stimuli, which is consistent with the left-right asymmetries in speech processing (Geschwind & Levitsky, 1968; Hickok & Poeppel, 2007).

We found more activated voxels for attending to audiovisual stimuli than for attending to either of the unisensory stimuli. This is consistent with previous evidence that modality-specific selective attention can reduce multisensory integration (Badde et al., 2020; Mozolic et al., 2008), though we cannot rule out the possibility that it is due to the difference in the number of studies. Notably, we found overlapping voxels between audiovisual attention with either visual or auditory attention, but there were no overlapping voxels between visual and auditory attention. This finding suggests that modality-specific attention not only possibly attenuates multisensory integration but also involve different neural activities (Chambers et al., 2004; Fairhall & Macaluso, 2009; Fernández et al., 2015). We found activation of the insula for auditory attention but not for visual attention. This difference can be interpreted by salience processing, in which the insula as a key node in the “salience network” has a central role in detecting behaviorally relevant signals (Menon & Uddin, 2010; Uddin, 2015).

Though there were highly divergent neural activities across contexts, we found that one brain region was consistently involved in audiovisual integration: superior temporal cortex. While this finding is consistent with the extensive prior literature suggesting the importance of superior temporal cortex in audiovisual integration (Beauchamp et al., 2004; Calvert & Thesen, 2004; Werner & Noppeney, 2010), our results highlight that there is not a singular integration region within the superior temporal cortex. Notably, there was no overlapping neural activity across levels for either analytical contrast, stimulus complexity or attention, but conjunction analyses showed that pairwise overlapping neural activity was in superior temporal cortex. Despite adjacent regions being activated under different contexts, the superior temporal cortex remains part of each of these distinct integration networks. It is possible that the superior temporal cortex is centrally located among different sites of audiovisual integration (visual cortex, subcortical node, and prefrontal cortex), which has the ease to get involved in all the possible neural pathways. Together, our findings highlight a flexible multiple pathways model for audiovisual integration, with superior temporal cortex as the central node in these neural assemblies.

Thus, our results support the idea that audiovisual integration can take place at a myriad of levels throughout the cortex and the identified areas partially depend on the choices made by researchers as to how they will induce and identify multisensory brain activity in their data. It provides support for the multiple pathways model in which audiovisual integration is associated with early cortical areas, subcortical areas, and higher association areas. The neural pathways for audiovisual integration appear to be a flexible rather than fixed network of brain regions with superior temporal cortex playing a central role. Together, these findings provide insights on the neural mechanisms of audiovisual integration.

## Acknowledgement

This work was supported by a Radboud Excellence Fellowship from Radboud University in Nijmegen, the Netherlands and funding from the University of South Carolina.

## Notes

Conflicts of Interest: None.

### Competing Interest Statement

The authors have declared no competing interest.

