## Supplementary Information for "Audiovisual Integration in the Human Brain: A Coordinate-based Meta-analysis"

**Supplementary Information for**
Audiovisual Integration in the Human Brain: A Coordinate-based Meta-analysis

Chuanji Gao*, Jessica J. Green, Xuan Yang, Sewon Oh, Jongwan Kim, and Svetlana V. Shinkareva*

Chuanji Gao

Svetlana V. Shinkareva


**This file includes:**

Supplementary text
Fig S1. Flowchart of the meta-analysis procedure.
Fig S2. Significant brain regions from the activation likelihood estimation (ALE) meta-analyses for audiovisual studies.
Table S1. Brain activation for different sensory modalities: visual-auditory and visual-tactile.
Table S2. Brain activation for different analytical contrasts.
Table S3. Brain activation for different analytical contrasts: shared voxels across contrasts.
Table S4. Brain activation for different stimuli: Simple, Complex-nonspeech and Complex-speech.
Table S5. Brain activation for different stimuli: shared voxels across stimulus types.
Table S6. Brain activation for different attention conditions: Audiovisual, Visual, Auditory and Passive.
Table S7. Brain activation for different attention conditions: shared voxels across attention conditions.

Legends for Datasets S1 to S2
SI References: 137 papers for all the studies included in the meta-analysis.

**Other supplementary materials for this manuscript include the following:**

Dataset S1. Summary of studies included for the meta-analysis of audiovisual integration.
Dataset S2. Summary of studies included for the meta-analysis of visual-tactile integration.

Supplementary Information Text

**Search terms**

Search terms included combinations of the three descriptors: one mandatory research question descriptor (“integration”), one research question descriptor (i.e., “multisensory”, “multi-sensory”, “multimodal”, “multi-modal”, “crossmodal”, “cross-modal”, “heteromodal”, “hetero-modal”, “polymodal”, “poly-modal”, “ploysensory”, “poly-sensory”, “intersensory”, “inter-sensory”, “audiovisual”, “audio-visual”, “audio AND visual”, “auditory-visual”, “visual-auditory”, “auditory AND visual”, “vision AND audition”, “sight AND sound”, “face AND voice”, “see AND hear”, “picture AND sound”, “visuotactile”, “visuo-tactile”, “visuo AND tactile”, “visuohaptic”, “visuo-haptic”, “visuo-somatosensory”, “visuo AND somatosensory”, “visual-tactile”, “visual AND tactile”, “visual-haptic”, “visual AND haptic”, “visual-somatosensory”, “visual AND somatosensory”, “vision AND touch”, “sight AND touch”, “see AND touch”, “visuo-thermal”, “visual-thermal”, “visuo AND thermal”, “visual AND thermal”, “audiotactile”, “audio-tactile”, “audio AND tactile”, “audio-haptic”, “audio AND haptic”, “audio-somatosensory”, “audio AND somatosensory”, “auditory-tactile”, “auditory AND tactile”, “auditory-haptic”, “auditory AND haptic”, “audition AND touch”, “hear AND touch”, “audio-thermal”, “auditory-thermal”, “audio AND thermal”, or “auditory AND thermal”) and one methodological descriptor (“fMRI”, “MRI”, “PET”, “BOLD”, “functional magnetic resonance imaging”, “functional imaging”, or “positron emission tomography”) for studies published until September 2020.

**Fig S1**. Flowchart of the meta-analysis procedure.

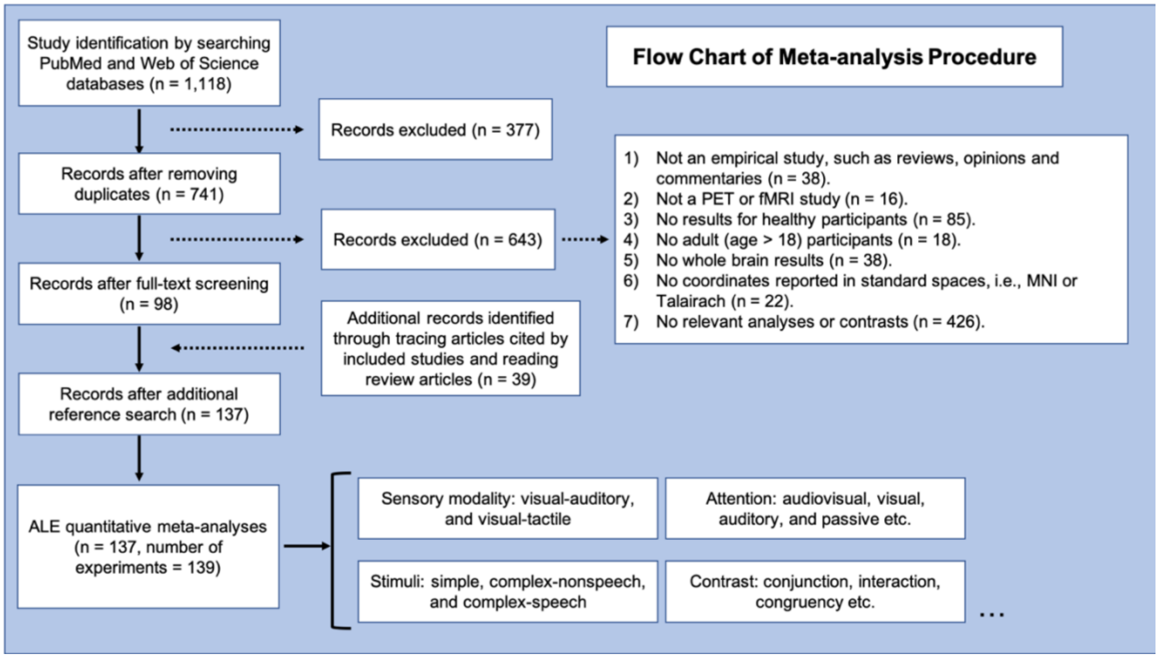

**Fig S2**. Significant brain regions from the activation likelihood estimation (ALE) meta-analyses for audiovisual studies. These findings showed that audiovisual integration is associated with multiple integration sites from early cortical areas, subcortical areas to higher association areas. Surface rendering was created using converted surface coordinates from MNI coordinates for visualization purpose.

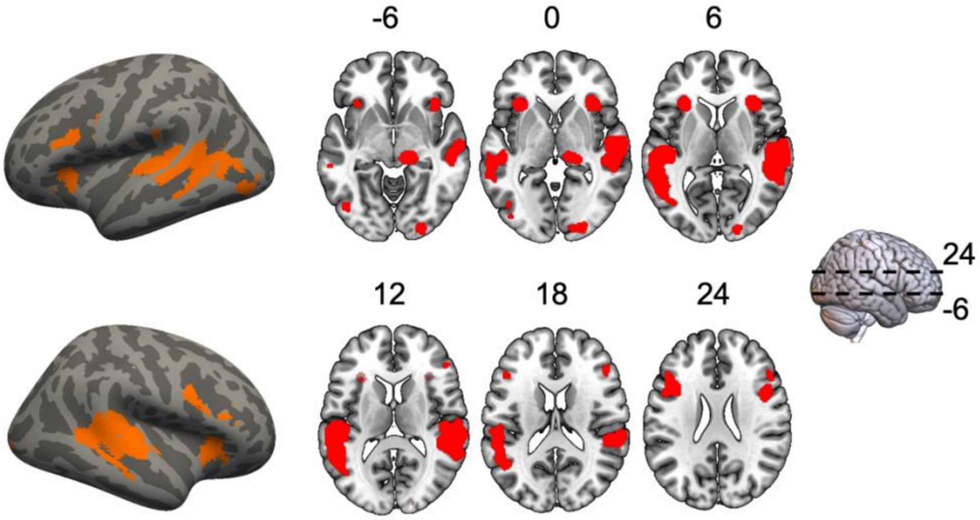

**Table S1**. Brain activation for different sensory modalities: visual-auditory and visual-tactile.

| Cluster Number. | Cluster Size ($mm^3$) | Anatomical Structure | MNI Coordinate | Z score |
|---|---|---|---|---|
| Visual-auditory | | | | |
| 1 | 17320 | L Superior Temporal Gyrus | -50 -22 8 | 8.19 |
| | | L Middle Temporal Gyrus | -56 -46 8 | 8.10 |
| | | L Fusiform Gyrus | -46 -72 -6 | 4.65 |
| | | L Middle Temporal Gyrus | -62 -34 -2 | 4.47 |
| | | L Middle Occipital Gyrus | -42 -68 8 | 3.86 |
| | | L Inferior Occipital Gyrus | -42 -84 2 | 3.60 |
| 2 | 16520 | R Superior Temporal Gyrus | 56 -34 10 | 7.24 |
| | | R Superior Temporal Gyrus | 60 -18 0 | 6.47 |
| | | R Superior Temporal Gyrus | 60 -48 10 | 5.53 |
| 3 | 3080 | R Inferior Frontal Gyrus | 48 10 28 | 5.56 |
| | | R Middle Frontal Gyrus | 48 32 20 | 3.79 |
| | | R Middle Frontal Gyrus | 50 22 30 | 3.64 |
| | | R Middle Frontal Gyrus | 50 34 14 | 3.40 |
| 4 | 2880 | L Middle Frontal Gyrus | -42 14 26 | 6.01 |
| | | L Middle Frontal Gyrus | -46 24 22 | 4.54 |
| | | L Precentral Gyrus | -46 4 34 | 3.50 |
| 5 | 2408 | R Insula | 34 24 4 | 7.61 |
| 6 | 1776 | L Insula | -30 24 6 | 6.15 |
| 7 | 1512 | R Thalamus | 14 -26 -4 | 5.73 |
| 8 | 1440 | L Medial Frontal Gyrus | -4 10 54 | 5.55 |
| | | L Superior Frontal Gyrus | -2 18 48 | 4.22 |
| 9 | 1288 | R Middle Occipital Gyrus | 24 -92 -4 | 4.45 |
| | | R Lingual Gyrus | 18 -94 6 | 4.03 |
| | | R Inferior Occipital Gyrus | 14 -94 -2 | 3.61 |
| Visual-tactile | | | | |
| 1 | 1208 | L Inferior Temporal Gyrus | -48 -68 4 | 4.78 |
| | | L Fusiform Gyrus | -46 -64 -8 | 3.96 |
| 2 | 1136 | R Inferior Parietal Lobule | 54 -30 48 | 5.02 |
| | | R Inferior Parietal Lobule | 42 -34 58 | 3.17 |
| 3 | 696 | L Inferior Parietal Lobule | -60 -26 34 | 4.69 |

L: left; R: right.

**Table S2**. Brain activation for different analytical contrasts.

| Cluster Number | Cluster Size ($mm^3$) | Anatomical Structure | MNI Coordinate | Z score |
|---|---|---|---|---|
| Conjunction | | | | |
| 1 | 7320 | R Superior Temporal Gyrus | 56 -34 10 | 6.65 |
| | | R Superior Temporal Gyrus | 58 -36 14 | 6.40 |
| | | R Superior Temporal Gyrus | 52 -42 8 | 6.32 |
| 2 | 4048 | L Superior Temporal Gyrus | -52 -48 12 | 6.09 |
| 3 | 2112 | R Thalamus | 14 -26 -4 | 7.31 |
| 4 | 1272 | L Thalamus | -14 -28 -4 | 5.50 |
| 5 | 840 | R Parahippocampal Gyrus | 26 -2 -16 | 4.39 |
| Interaction | | | | |
| 1 | 2496 | L Middle Temporal Gyrus | -62 -32 2 | 4.47 |
| | | L Middle Temporal Gyrus | -58 -40 6 | 4.16 |
| | | L Superior Temporal Gyrus | -48 -24 8 | 3.91 |
| | | L Superior Temporal Gyrus | -52 -28 14 | 3.79 |
| 2 | 2336 | R Superior Temporal Gyrus | 52 -18 2 | 4.37 |
| | | R Superior Temporal Gyrus | 62 -24 4 | 4.36 |
| | | R Postcentral Gyrus | 60 -16 14 | 3.80 |
| Congruent > Incongruent | | | | |
| 1 | 3184 | R Superior Temporal Gyrus | 62 -14 -4 | 4.62 |
| | | R Superior Temporal Gyrus | 52 -22 6 | 4.34 |
| | | R Superior Temporal Gyrus | 52 -26 -6 | 3.91 |
| 2 | 1528 | L Inferior Temporal Gyrus | -46 -72 -2 | 4.55 |
| | | L Inferior Occipital Gyrus | -42 -84 2 | 4.27 |
| | | L Middle Occipital Gyrus | -40 -72 8 | 3.78 |
| 3 | 1448 | L Superior Temporal Gyrus | -52 -22 6 | 5.04 |
| Incongruent > Congruent | | | | |
| 1 | 1800 | L Middle Frontal Gyrus | -42 14 26 | 5.37 |
| 2 | 1752 | L Superior Frontal Gyrus | 0 22 48 | 5.20 |
| 3 | 1208 | R Inferior Frontal Gyrus | 44 10 26 | 4.29 |
| | | R Inferior Frontal Gyrus | 50 18 20 | 3.88 |
| 4 | 992 | R Superior Temporal Gyrus | 56 -32 2 | 4.67 |
| 5 | 952 | L Middle Temporal Gyrus | -56 -44 6 | 4.14 |

L: left; R: right.

**Table S3.** Brain activation for different analytical contrasts: shared voxels across contrasts.

| Conjunction | Voxel counts | Center of mass | Brain region |
|---|---|---|---|
| Conjunction ∩ Interaction | 21 voxels | [60 -31 9] | R. Superior Temporal Cortex |
| | 73 voxels | [-56 -45 10] | L. Middle Temporal Cortex |
| Conjunction ∩ Congruency | 4 voxels | [53 -28 -2] | R. Superior Temporal Cortex |
| Interaction ∩ Congruency | 73 voxels | [55 -20 2] | R. Superior Temporal Cortex |
| | 23 voxels | [-49 -23 8] | L. Superior Temporal Cortex |

Note. The conjunction between contrasts were performed using the conservative minimum statistic (Nichols et al., 2005) in which only voxels significant on a corrected level in each contrast were considered. Anatomical location labeling is based on the AAL3 atlas (http://www.gin.cnrs.fr/en/tools/aal/).

**Table S4**. Brain activation for different stimuli: Simple, Complex-nonspeech and Complex-speech.

| Cluster Number | Cluster Size ($mm^3$) | Anatomical Structure | MNI Coordinate | Z score |
|---|---|---|---|---|
| Simple | | | | |
| 1 | 2280 | L Superior Temporal Gyrus | -52 -22 6 | 5.32 |
| | | L Superior Temporal Gyrus | -44 -34 14 | 4.01 |
| 2 | 1976 | R Transverse Temporal Gyrus | 50 -24 10 | 5.30 |
| | | R Superior Temporal Gyrus | 56 -34 14 | 4.14 |
| 3 | 1416 | R Inferior Frontal Gyrus | 48 10 30 | 5.34 |
| | | R Middle Frontal Gyrus | 50 22 32 | 4.08 |
| Complex-nonspeech | | | | |
| 1 | 1616 | R Superior Temporal Gyrus | 60 -18 -6 | 4.46 |
| | | R Superior Temporal Gyrus | 58 -10 0 | 4.05 |
| Complex-speech | | | | |
| 1 | 13992 | L Superior Temporal Gyrus | -54 -52 10 | 7.57 |
| | | L Middle Temporal Gyrus | -56 -46 8 | 7.51 |
| | | L Superior Temporal Gyrus | -62 -30 10 | 4.96 |
| | | L Superior Temporal Gyrus | -50 -22 8 | 4.85 |
| | | L Superior Temporal Gyrus | -50 -30 2 | 3.72 |
| 2 | 10144 | R Superior Temporal Gyrus | 52 -40 8 | 7.28 |
| | | R Superior Temporal Gyrus | 54 -32 4 | 7.21 |
| | | R Superior Temporal Gyrus | 52 -14 2 | 3.90 |
| 3 | 3040 | L Inferior Frontal Gyrus | -40 12 26 | 5.77 |
| | | L Middle Frontal Gyrus | -46 24 22 | 4.65 |
| 4 | 1608 | R Middle Occipital Gyrus | 24 -92 -4 | 4.69 |
| | | R Lingual Gyrus | 18 -96 6 | 4.51 |
| | | R Lingual Gyrus | 6 -92 -4 | 3.73 |
| | | R Inferior Occipital Gyrus | 14 -92 -2 | 3.59 |
| 5 | 1512 | R Thalamus | 14 -28 -4 | 5.52 |
| 6 | 1208 | L Medial Frontal Gyrus | -2 10 56 | 4.63 |
| | | L Superior Frontal Gyrus | 0 22 48 | 4.15 |

L: left; R: right.

**Table S5.** Brain activation for different stimuli: shared voxels across stimulus types.

| Conjunction | Voxel counts | Center of mass | Brain region |
|---|---|---|---|
| Simple ∩ Complex-speech | 126 voxels | [54 -28 9] | R. Superior Temporal Cortex |
| | 181 voxels | [-52 -24 9] | L. Superior Temporal Cortex |
| Complex-nonspeech ∩ Complex-speech | 63 voxels | [60 -16 -2] | R. Superior Temporal Cortex |

Note. The conjunction between analyses were performed using the conservative minimum statistic (Nichols et al., 2005) in which only voxels significant on a corrected level in each analysis were considered. Anatomical location labeling is based on the AAL3 atlas (http://www.gin.cnrs.fr/en/tools/aal/).

**Table S6**. Brain activation for different attention conditions: Audiovisual, Visual, Auditory and Passive.

| Cluster Number | Cluster Size ($mm^3$) | Anatomical Structure | MNI Coordinate | Z score |
|---|---|---|---|---|
| Audiovisual | | | | |
| 1 | 8096 | R Insula | 50 -14 2 | 5.09 |
| | | R Middle Temporal Gyrus | 60 -44 10 | 5.01 |
| | | R Middle Temporal Gyrus | 54 -42 8 | 4.85 |
| | | R Superior Temporal Gyrus | 60 -16 -6 | 4.84 |
| | | R Superior Temporal Gyrus | 64 -32 14 | 3.96 |
| | | R Superior Temporal Gyrus | 50 -30 0 | 3.89 |
| 2 | 8096 | L Superior Temporal Gyrus | -58 -44 8 | |
| | | L Superior Temporal Gyrus | -48 -24 8 | |
| | | L Middle Temporal Gyrus | -54 -56 8 | |
| | | L Middle Temporal Gyrus | -62 -34 -2 | |
| | | L Superior Temporal Gyrus | -54 -54 16 | |
| | | L Superior Temporal Gyrus | -44 -58 20 | |
| | | L Superior Temporal Gyrus | -48 -26 -2 | |
| | | L Superior Temporal Gyrus | -66 -22 0 | |
| 3 | 1680 | R Thalamus | 12 -28 -4 | |
| 4 | 1608 | R Insula | 34 24 4 | |
| 5 | 1312 | L Inferior Frontal Gyrus | -46 10 28 | |
| | | L Precentral Gyrus | -46 4 34 | |
| Visual | | | | |
| 1 | 1856 | R Superior Temporal Gyrus | 54 -34 12 | 4.67 |
| | | R Superior Temporal Gyrus | 60 -48 12 | 4.38 |
| 2 | 896 | R Superior Temporal Gyrus | 60 -10 2 | 4.53 |
| Auditory | | | | |
| 1 | 1832 | L Superior Temporal Gyrus | -44 -34 14 | 4.31 |
| | | L Superior Temporal Gyrus | -46 -30 10 | 3.88 |
| 2 | 1824 | L Medial Frontal Gyrus | -2 8 56 | 5.15 |
| | | L Medial Frontal Gyrus | -2 16 46 | 4.45 |
| 3 | 1568 | L Insula | -30 22 6 | 5.34 |
| | | L Insula | -34 22 0 | 5.03 |
| 4 | 1448 | R Insula | 34 26 6 | 5.13 |
| Passive | | | | |
| 1 | 5920 | L Superior Temporal Gyrus | -54 -50 10 | 6.23 |
| | | L Middle Temporal Gyrus | -50 -62 8 | 5.06 |
| 2 | 4888 | R Middle Temporal Gyrus | 58 -34 8 | 5.59 |
| | | R Superior Temporal Gyrus | 52 -40 8 | 5.06 |
| | | R Superior Temporal Gyrus | 58 -20 2 | 4.87 |
| 3 | 864 | L Middle Frontal Gyrus | -44 18 30 | 4.33 |

| | | | | |
|---|---|---|---|---|
| 4 | 816 | R Middle Occipital Gyrus | 28 -88 -4 | 4.02 |
| | | R Lingual Gyrus | 22 -90 4 | 3.57 |

L: left; R: right.

**Table S7.** Brain activation for different attention conditions: shared voxels across attention conditions.

| Contrast | Voxel counts | Center of mass | Brain region |
|---|---|---|---|
| Audiovisual ∩ Visual | 34 voxels | [62 -13 0] | R. Superior Temporal Cortex |
| | 167 voxels | [56 -41 11] | R. Superior Temporal Cortex |
| Audiovisual ∩ Auditory | 111 voxels | [36 24 3] | R. Insula |
| | 87 voxels | [-51 -27 9] | L. Superior Temporal Cortex |

Note. The conjunction between analyses were performed using the conservative minimum statistic (Nichols et al., 2005) in which only voxels significant on a corrected level in each analysis were considered. Anatomical location labeling is based on the AAL3 atlas (http://www.gin.cnrs.fr/en/tools/aal/).

**Dataset S1 (separate file).** Summary of studies included for the meta-analysis of audiovisual integration.

**Dataset S2 (separate file).** Summary of studies included for the meta-analysis of visual-tactile integration.

**SI References**
**These are the 137 papers for all the studies included in the meta-analysis.**

69. Watson, R., Latinus, M., Noguchi, T., Garrod, O., Crabbe, F., & Belin, P. (2014). Crossmodal Adaptation in Right Posterior Superior Temporal Sulcus during Face-Voice Emotional Integration. The Journal of Neuroscience, 34(20), 6813–6821. https://doi.org/10.1523/jneurosci.4478-13.2014
70. Vander Wyk, B. C., Ramsay, G. J., Hudac, C. M., Jones, W., Lin, D., Klin, A., Lee, S. M., & Pelphrey, K. A. (2010). Cortical integration of audio-visual speech and non-speech stimuli. Brain Cogn, 74(2), 97–106. https://doi.org/10.1016/j.bandc.2010.07.002
71. Hadjikhani, N., & Roland, P. E. (1998). Cross-modal transfer of information between the tactile and the visual representations in the human brain: A positron emission tomographic study. Journal of Neuroscience, 18(3), 1072–1084. https://doi.org/10.1523/jneurosci.18-03-01072.1998
72. Rosemann, S., Smith, D., Dewenter, M., & Thiel, C. M. (2020). Age-related hearing loss influences functional connectivity of auditory cortex for the McGurk illusion. Cortex, 129, 266–280. https://doi.org/10.1016/j.cortex.2020.04.022
73. Pourtois, G., de Gelder, B., Bol, A., & Crommelinck, M. (2005). Perception of facial expressions and voices and of their combination in the human brain. Cortex, 41(1), 49–59. https://doi.org/10.1016/s0010-9452(08)70177-1
74. Kim, H., Hahm, J., Lee, H., Kang, E., Kang, H., & Lee, D. S. (2015). Brain Networks Engaged in Audiovisual Integration During Speech Perception Revealed by Persistent Homology-Based Network Filtration. Brain Connectivity, 5(4), 245–258. https://doi.org/10.1089/brain.2013.0218
75. Saito, D. N., Okada, T., Morita, Y., Yonekura, Y., & Sadato, N. (2003). Tactile-visual cross-modal shape matching: a functional MRI study. Brain Res Cogn Brain Res, 17(1), 14–25. https://doi.org/10.1016/s0926-6410(03)00076-4
76. von Saldern, S., & Noppeney, U. (2013). Sensory and Striatal Areas Integrate Auditory and Visual Signals into Behavioral Benefits during Motion Discrimination. Journal of Neuroscience, 33(20), 8841–8849. https://doi.org/10.1523/jneurosci.3020-12.2013
77. Meehan, S. K., & Staines, W. R. (2009). Task-relevance and temporal synchrony between tactile and visual stimuli modulates cortical activity and motor performance during sensory-guided movement. Hum Brain Mapp, 30(2), 484–496. https://doi.org/10.1002/hbm.20520
78. Lamichhane, B., & Dhamala, M. (2015). The Salience Network and Its Functional Architecture in a Perceptual Decision: An Effective Connectivity Study. Brain Connect, 5(6), 362–370. https://doi.org/10.1089/brain.2014.0282
79. Holle, H., Obleser, J., Rueschemeyer, S. A., & Gunter, T. C. (2010). Integration of iconic gestures and speech in left superior temporal areas boosts speech comprehension under adverse listening conditions. Neuroimage, 49(1), 875–884. https://doi.org/10.1016/j.neuroimage.2009.08.058
80. Guediche, S., Zhu, Y., Minicucci, D., & Blumstein, S. E. (2019). Written sentence context effects on acoustic-phonetic perception: fMRI reveals cross-modal semantic-perceptual interactions. Brain and Language, 199. https://doi.org/10.1016/j.bandl.2019.104698
81. Ehrsson, H. H., Spence, C., & Passingham, R. E. (2004). That’s my hand! Activity in premotor cortex reflects feeling of ownership of a limb. Science, 305(5685), 875–877. https://doi.org/10.1126/science.1097011
82. Doehrmann, O., Weigelt, S., Altmann, C. F., Kaiser, J., & Naumer, M. J. (2010). Audiovisual Functional Magnetic Resonance Imaging Adaptation Reveals Multisensory Integration Effects in Object-Related Sensory Cortices. Journal of Neuroscience, 30(9), 3370–3379. https://doi.org/10.1523/jneurosci.5074-09.2010
83. Noesselt, T., Rieger, J. W., Schoenfeld, M. A., Kanowski, M., Hinrichs, H., Heinze, H. J., & Driver, J. (2007). Audiovisual temporal correspondence modulates human multisensory superior temporal sulcus plus primary sensory cortices. Journal of Neuroscience, 27(42), 11431–11441. https://doi.org/10.1523/JNEUROSCI.2252-07.2007
84. Benoit, M. M. K., Raij, T., Lin, F. H., Jääskeläinen, I. P., & Stufflebeam, S. (2010). Primary and multisensory cortical activity is correlated with audiovisual percepts. Human Brain Mapping, 31(4), 526–538. https://doi.org/10.1002/hbm.20884
85. Biau, E., Morís Fernández, L., Holle, H., Avila, C., & Soto-Faraco, S. (2016). Hand gestures as visual prosody: BOLD responses to audio-visual alignment are modulated by the communicative nature of the stimuli. NeuroImage, 132, 129–137. https://doi.org/10.1016/j.neuroimage.2016.02.018
86. Naghavi, H. R., Eriksson, J., Larsson, A., & Nyberg, L. (2011). Cortical regions underlying successful encoding of semantically congruent and incongruent associations between common

auditory and visual objects. Neuroscience Letters, 505(2), 191–195. https://doi.org/10.1016/j.neulet.2011.10.022
87. Guterstam, A., Gentile, G., & Ehrsson, H. H. (2013). The Invisible Hand Illusion: Multisensory Integration Leads to the Embodiment of a Discrete Volume of Empty Space. Journal of Cognitive Neuroscience, 25(7), 1078–1099. https://doi.org/10.1162/jocn_a_00393
88. Werner, S., & Noppeney, U. (2009). Superadditive Responses in Superior Temporal Sulcus Predict Audiovisual Benefits in Object Categorization. Cerebral Cortex, 20(8), 1829–1842. https://doi.org/10.1093/cercor/bhp248
89. van Atteveldt, N. M., Blau, V. C., Blomert, L., & Goebel, R. (2010). fMR-adaptation indicates selectivity to audiovisual content congruency in distributed clusters in human superior temporal cortex. BMC Neuroscience, 11. https://doi.org/10.1186/1471-2202-11-11
90. Bushara, K. O., Hanakawa, T., Immisch, I., Toma, K., Kansaku, K., & Hallett, M. (2003). Neural correlates of cross-modal binding. Nature Neuroscience, 6(2), 190–195. https://doi.org/10.1038/nn993
91. Lu, L., & Liu, B. L. (2020). Revealing the multisensory modulation of auditory stimulus in degraded visual object recognition by dynamic causal modeling. Brain Imaging and Behavior, 14(4), 1187–1198. https://doi.org/10.1007/s11682-019-00134-3
92. Noesselt, T., Bergmann, D., Heinze, H. J., Münte, T., & Spence, C. (2012). Coding of multisensory temporal patterns in human superior temporal sulcus. Frontiers in Integrative Neuroscience, 6(AUGUST 2012), 1–14. https://doi.org/10.3389/fnint.2012.00064
93. Park, J. Y., Gu, B. M., Kang, D. H., Shin, Y. W., Choi, C. H., Lee, J. M., & Kwon, J. S. (2010). Integration of cross-modal emotional information in the human brain: An fMRI study. Cortex, 46(2), 161–169. https://doi.org/10.1016/j.cortex.2008.06.008
94. Luttke, C. S., Ekman, M., van Gerven, M. A. J., & de Lange, F. P. (2016). Preference for Audiovisual Speech Congruency in Superior Temporal Cortex. Journal of Cognitive Neuroscience, 28(1), 1–7. https://doi.org/10.1162/jocn_a_00874
95. Watkins, S., Shams, L., Tanaka, S., Haynes, J. D., & Rees, G. (2006). Sound alters activity in human V1 in association with illusory visual perception. NeuroImage, 31(3), 1247–1256. https://doi.org/10.1016/j.neuroimage.2006.01.016
96. Morís Fernández, L., Visser, M., Ventura-Campos, N., Ávila, C., Soto-Faraco, S., Fernandez, L. M., Visser, M., Ventura-Campos, N., Avila, C., & Soto-Faraco, S. (2015). Top-down attention regulates the neural expression of audiovisual integration. NeuroImage, 119, 272–285. https://doi.org/10.1016/j.neuroimage.2015.06.052
97. Gentile, G., Björnsdotter, M., Petkova, V. I., Abdulkarim, Z., & Ehrsson, H. H. (2015). Patterns of neural activity in the human ventral premotor cortex reflect a whole-body multisensory percept. Neuroimage, 109, 328–340. https://doi.org/10.1016/j.neuroimage.2015.01.008
98. Laing, M., Rees, A., & Vuong, Q. C. (2015). Amplitude-modulated stimuli reveal auditory-visual interactions in brain activity and brain connectivity. Frontiers in Psychology, 6. https://doi.org/10.3389/fpsyg.2015.01440
99. Noppeney, U., Ostwald, D., & Werner, S. (2010). Perceptual decisions formed by accumulation of audiovisual evidence in prefrontal cortex. Journal of Neuroscience, 30(21), 7434–7446. https://doi.org/10.1523/JNEUROSCI.0455-10.2010
100. Kang, E., Lee, D. S., Kang, H., Hwang, C. H., Oh, S. H., Kim, C. S., Chung, J. K., & Lee, M. C. (2006). The neural correlates of cross-modal interaction in speech perception during a semantic decision task on sentences: a PET study. Neuroimage, 32(1), 423–431. https://doi.org/10.1016/j.neuroimage.2006.03.016
101. Brozzoli, C., Gentile, G., & Henrik Ehrsson, H. (2012). That’s near my hand! Parietal and premotor coding of hand-centered space contributes to localization and self-attribution of the hand. Journal of Neuroscience, 32(42), 14573–14582. https://doi.org/10.1523/JNEUROSCI.2660-12.2012
102. Gentile, G., Guterstam, A., Brozzoli, C., & Ehrsson, H. H. (2013). Disintegration of Multisensory Signals from the Real Hand Reduces Default Limb Self-Attribution: An fMRI Study. Journal of Neuroscience, 33(33), 13350–13366. https://doi.org/10.1523/jneurosci.1363-13.2013
103. Naghavi, H. R., Eriksson, J., Larsson, A., & Nyberg, L. (2007). The claustrum/insula region integrates conceptually related sounds and pictures. Neurosci Lett, 422(1), 77–80. https://doi.org/10.1016/j.neulet.2007.06.009
